# Antiviral activity of plant juices and green tea against SARS-CoV-2 and influenza virus *in vitro*

**DOI:** 10.1101/2020.10.30.360545

**Authors:** Bruno Frank, Carina Conzelmann, Tatjana Weil, Rüdiger Groß, Peggy Jungke, Maren Eggers, Janis A. Müller, Jan Münch, Uwe Kessler

## Abstract

Many plant juices, extracts and teas have been shown to possess antiviral activity. We here analyzed the virucidal activity of black chokeberry *(Aronia melanocarpa),* pomegranate *(Punica granatum),* and elderberry *(Sambucus nigra)* juice, as well as green tea *(Camellia sinensis)* against different respiratory viruses. We found that all tested plant derived products effectively inactivated influenza virus, whereas only chokeberry juice diminished SARS-CoV-2 and vaccinia virus infectivity. None of the products inactivated non-enveloped human adenovirus type 5. Thus, black chokeberry juice exerts virucidal activity against different enveloped viral pathogens under *in vitro* conditions. Whether application of virucidal juices or green tea as oral rinses may lower viral loads in the oral cavity *in vivo* remains to be evaluated.

## Background

Respiratory viruses such as influenza viruses and coronaviruses pose a significant threat to global health and are a substantial social, economic and healthcare burden, recently exemplified by the coronavirus disease 2019 (COVID-19) pandemic caused by severe acute respiratory syndrome coronavirus 2 (SARS-CoV-2) (1). For SARS-CoV-2, the long incubation period of up to 14 days, subclinical course and high transmissibility before onset of symptoms has led to unprecedented spread around the globe (1, 2). Respiratory viruses initially infect the upper airways, both the naso- and oropharyngeal areas, where they amplify, cause respiratory symptoms (3) and spread to new hosts. Recent studies suggest gargling with commercial oral rinses may reduce virus spread and potentially infection (4, 5). Several natural products also have direct antiviral activity or may ameliorate symptoms of respiratory infections. Pomegranate *(Punica granatum)* (6) and black chokeberry *(Aronia melanocarpa)* (7) extracts have been shown to be antivirally active against influenza viruses *in vitro,* elderberry syrup *(Sambucus nigra)* has shown improved symptom relief in influenza patients (8), and a meta-analysis showed that gargling green tea *(Camellia sinensis)* lowered the incidence of influenza infections (9). Natural products with a broad-spectrum antiviral activity would therefore be useful to reduce spread of respiratory viruses in the population, as they are inexpensive and rapidly deployable. Here, we evaluated the *in vitro* virucidal activity of green tea and herbal juices with prospective use as oral rinses against the enveloped respiratory viruses, SARS-CoV-2 and influenza A virus (IAV) and the naked adenovirus type 5 (AdV5). We found that IAV is highly susceptible to inactivation by all tested substances. SARS-CoV-2 was less affected, however, inactivated by chokeberry juice and sensitive to green tea and pomegranate juice. AdV5 was resistant to most products, but viral titers were slightly reduced by chokeberry juice. These findings underline the potential of common plant derived food products to inactivate enveloped viruses *in vitro,* where chokeberry juice was the most potent natural product tested herein. Whether chokeberry juice applied as oral rinse may lower viral loads in the oral cavity *in vivo,* remains to be assessed.

## Methods

### Herbal substances

Green tea (Bio-Grüntee Japan Sencha Tee-Gschwendner Nr. 700; pH 4.46) was prepared by infusing 3 g of leaves with 0.1 g ascorbic acid (Sigma Aldrich) for 4 min in 300 ml freshly boiled water under gentle movement, followed by filtration. Black chokeberry juice (Bio-Aronia Direktsaft Fa. Aronia original L2719; pH: 3.69), pomegranate juice (Satower Granatapfelsaft Direktsaft klar; pH: 2.99), and elderberry juice (Satower Fliederbeersaft; pH: 4.13) with valid best before date were kept refrigerated until use.

### Cell culture

Vero E6 *(Cercopithecus aethiops* derived epithelial kidney) cells were grown in Dulbecco’s modified Eagle’s medium (DMEM, Gibco) which was supplemented with 2.5% heat-inactivated fetal calf serum (FCS), 100 units/ml penicillin, 100 μg/ml streptomycin, 2 mM L-glutamine, 1 mM sodium pyruvate, and 1x non-essential amino acids. Madin Darby canine kidney cells (MDCK) and A549 (adenocarcinomic human alveolar basal epithelial) cells were grown in minimal essential medium with Earle’s salts (EMEM; Biochrom AG, Berlin, Germany) supplemented with 1% non-essential aminoacids (Biochrom AG, Berlin, Germany), 10% FCS. BHK-21 *(Mesocricetus auratus* kidney) cells were grown in DMEM (CCPro) with 10% FCS. For experiments, cells were seeded in medium containing 2% FCS. Cells were incubated at 37°C in a 5% CO_2_ humidified incubator.

### Virus test strains and cultivation

Virus was propagated by inoculation of respective target cells and culturing until strong cytopathic effect was visible. The supernatant was then harvested, centrifuged to deplete cellular debris, aliquoted and stored at −80°C as virus stocks. Modified vaccinia virus Ankara (provided from the Institute of Animal Hygiene and Veterinary Public Health of the University Leipzig) was passaged on BHK-21 cells (provided by Friedrich Löffler institute), influenza A virus A/H1N1/Brisbane/59/2007 (Novartis Vaccines and Diagnostics GmbH & Co. KG) on MDCK cells (ATCC), adenovirus type 5, strain adenoid 75 (kindly provided by Prof. Sauerbrei, University of Jena, Jena, Germany) on A549 cells (ATCC) and SARS-CoV-2 BetaCoV/France/IDF0372/2020 (obtained through European Virus Archive global) on Vero E6 cells (ATCC).

### Infection assays

To determine the virucidal activity of the herbal substances, they were mixed with the respective virus, incubated for a specified contact time at room temperature, and the remaining infectivity determined by tissue culture infectious dose 50 (TCID_50_) endpoint titration. This quantitative suspension test as described in EN 14476 (10) was performed for Modified vaccinia virus Ankara (MVA), influenza A virus (IAV) and adenovirus type 5 (AdV5) by incubating the respective virus with chokeberry, elderberry, or pomegranate juice, green tea, or buffer as control. Briefly, the efficacy of the tested products was examined as an 80% solution in the presence of 10% interfering substance (5% (w/v) BSA Fraction V (Sigma Aldrich), 0.4% (w/v) Mucin bovine Glandula submandibularis Type I-S (Sigma Aldrich), 5% (w/v) yeast extract (Sigma Aldrich)). SARS-CoV-2 was analyzed in 90% product. After the specified contact time, the test mixture was serially diluted 10-fold and titrated onto a 96 microtiter plate containing a confluent monolayer of the respective target cells in sextuplicates and the cells cultured until strong CPE was visible. IAV infected cells were additionally immunostained. Cells were then examined with a light microscope, the infected wells counted, and TCID_50_ calculated according to Spearman-Kaerber. If the cytotoxicity of the compound succeeded the lower limit of quantification (LLOQ), the titer was adjusted accordingly. The virucidal activity was determined by the difference of the logarithmic titer of the virus control minus the logarithmic titer of the virus incubated with the substance to test.

## Results

To assess the virucidal potential of four plant-derived products, we performed a quantitative suspension test using MVA (EN 14476 (10)) which is a resilient surrogate virus that is used for the validation of virucidal disinfectants against all enveloped viruses according to the European Guidance on the Biocidal Products Regulation (11). While no reduction in viral titer was observed upon incubation with control buffer, 5-minute incubation with chokeberry juice, elderberry juice, pomegranate juice, or green tea yielded a 3.17, 0.67, 1.0 or 1.0 log_10_ decrease in infectivity, respectively (Figure 1, Table 1), indicating that the tested products are generally active against enveloped viruses. An incubation time of 20 minutes was only marginally more potent, suggesting a rapid acting antiviral effect. We then analyzed the two respiratory enveloped viruses responsible for the “swine flu” in 2009/2010 and the ongoing COVID-19 pandemic, IAV and SARS-CoV-2, respectively, as well as AdV5 as a naked control virus. A 5-minute incubation with chokeberry juice yielded most potent antiviral activities and inactivated IAV, SARS-CoV-2 and also AdV5 to 99.99%, 96.98%, and 93.23%, respectively (Figure 1, Table 1). IAV was most susceptible to all products and infectivity reduced >99% by elderberry juice, pomegranate juice and green tea. SARS-CoV-2 titers were reduced approximately 80% by pomegranate juice and green tea already after 1-minute incubation, however, unaffected by elderberry juice, corresponding to the results obtained with the more resistant surrogate MVA. The naked AdV5 was resistant to three out of four products, however, susceptible to chokeberry juice (Figure 1, Table 1). In summary, IAV is highly susceptible to all analyzed products, whereas SARS-CoV-2 can be efficiently inactivated by chokeberry juice and is to a lower level affected by pomegranate juice or green tea.

**Figure 1:**
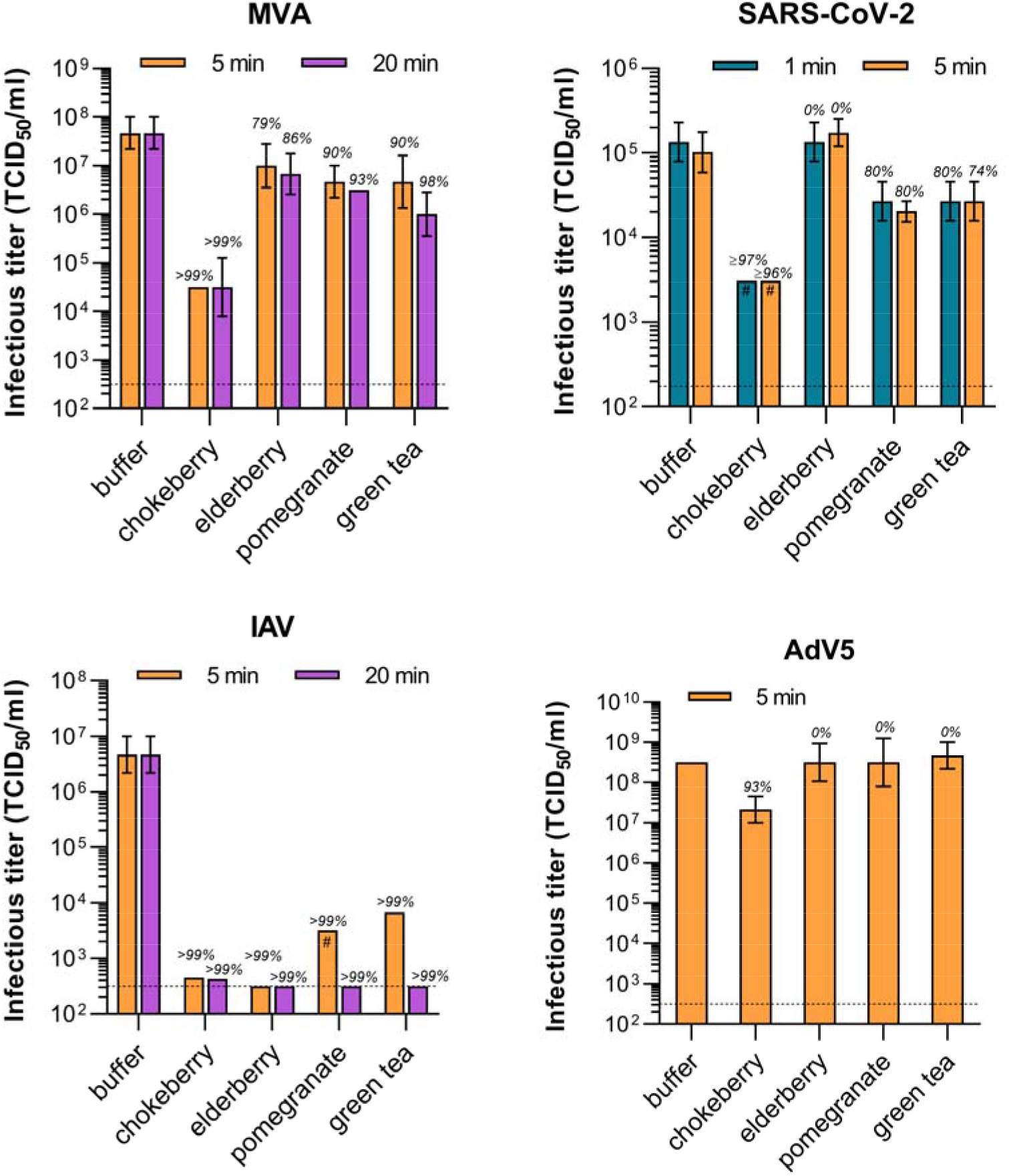
Virucidal activity of natural products against MVA, IAV, SARS-CoV-2 and AdV5. MVA, IAV (A/H1N1/Brisbane/59/2007), SARS-CoV-2 (BetaCoV/France/IDF0372/2020), or AdV5 (adenoid 75) were incubated with the plant derived products for indicated contact times before serial titration and inoculation of target cells. Viral titers were determined by monitoring cytopathic effect and calculated as tissue culture infectious dose 50 (TCID_50_) according to Spearman-Kaerber. The lower limit of quantification (LLOQ) is defined by limit of titration (dotted line) or cytotoxicity of the compound (#). Error bars indicate standard deviation and italics above corresponding bars the decrease of titers compared to control.

**Table 1:** Antiviral activity of natural products against MVA, IAV, SARS-CoV-2 and AdV5. Log10 reduction factor and antiviral activity of chokeberry, elderberry, pomegranate juice and green tea against MVA, IAV, SARS-CoV-2 and AdV5 after indicated contact times. MVA, Modified vaccinia virus Ankara; IAV, influenza A virus (A/H1N1/Brisbane/59/2007); SARS-CoV-2, severe acute respiratory syndrome coronavirus 2 (BetaCoV/France/IDF0372/2020); AdV5 adenovirus type 5 (adenoid 75).

| virus | contact time (min) |  | control | chokeberry | elderberry | pomegranate | green tea |
| --- | --- | --- | --- | --- | --- | --- | --- |
| **MVA** | **5** | titer | 7.67 ± 0.33 | 4.50 ± 0.00 | 7.00 ± 0.45 | 6.67 ± 0.33 | 6.67 ± 0.54 |
|  |  | log<sub>10</sub> reduction factor |  | 3.17 | 0.67 | 1.00 | 1.00 |
|  |  | infectivity reduction (%) |  | 99.93 | 78.62 | 90.00 | 90.00 |
|  | **20** | titer | 7.67 ± 0.33 | 4.50 ± 0.60 | 6.83 ± 0.42 | 6.50 ± 0.00 | 6.00 ± 0.45 |
|  |  | log<sub>10</sub> reduction factor |  | 3.17 | 0.84 | 1.17 | 1.67 |
|  |  | infectivity reduction (%) |  | 99.93 | 85.55 | 93.24 | 97.86 |
| **IAV** | **5** | titer | 6.67 ± 0.33 | 2.66 ± 0.0 | ≤ 2.5 ± 0.0 | ≤ 3.5 ± 0.0 | 3.83 ± 0.0 |
|  |  | log<sub>10</sub> reduction factor |  | 4.01 | ≥ 4.17 | ≥ 3.17 | 2.84 |
|  |  | infectivity reduction (%) |  | 99.99 | ≥ 99.99 | ≥ 99.93 | 99.86 |
|  | **20** | titer | 6.67 ± 0.33 | 2.63 ± 0.0 | ≤ 2.5 ± 0.0 | ≤ 2.5 ± 0.0 | ≤ 2.5 ± 0.0 |
|  |  | log<sub>10</sub> reduction factor |  | 4.04 | ≥ 4.17 | ≥ 4.17 | ≥ 4.17 |
|  |  | infectivity reduction (%) |  | 99.99 | ≥ 99.99 | ≥ 99.99 | ≥ 99.99 |
| **SARS-CoV-2** | **1** | titer | 5.13 ± 0.23 | ≤ 3.49 ± 0.00 | 5.13 ± 0.23 | 4.43 ± 0.23 | 4.43 ± 0.23 |
|  |  | log<sub>10</sub> reduction factor |  | ≥ 1.64 | 0 | 0.70 | 0.70 |
|  |  | infectivity reduction (%) |  | ≥ 97.71 | 0 | 80.05 | 80.05 |
|  | **5** | titer | 5.01 ± 0.24 | ≤ 3.49 ± 0 | 5.24 ± 0.16 | 4.31 ± 0.12 | 4.43 ± 0.23 |
|  |  | log<sub>10</sub> reduction factor |  | ≥ 1.52 | 0 | 0.70 | 0.58 |
|  |  | infectivity reduction (%) |  | ≥ 96.98 | 0 | 80.05 | 73.70 |
| **AdV5** | **5** | titer | 8.50 ± 0.00 | 7.33 ± 0.33 | 8.50 ± 0.47 | 8.50 ± 0.60 | 8.67 ± 0.33 |
|  |  | log<sub>10</sub> reduction factor |  | 1.17 | 0 | 0 | 0 |
|  |  | infectivity reduction (%) |  | 93.24 | 0 | 0 | 0 |

## Discussion

We examined and compared the virucidal activities of four natural beverages on a surrogate and three respiratory viruses and found that chokeberry juice, green tea and pomegranate juice reduced infectious titers of enveloped viruses with chokeberry juice being most efficient. The tested food products showed the highest antiviral efficacy against IAV, with ≥ 4 log_10_, which corresponds to the effectiveness of typical disinfectants. The high susceptibility of IAV (H1N1), which is also a representative of influenza B and other influenza strains with regard to chemical stability, indicates low resilience of this virus family. SARS-CoV-2 behaved similar to the European model virus for enveloped viruses, MVA, and proved to be more resilient. Notably, activity against MVA is prerequisite for validation of general disinfectant property of chemicals according to the European Chemicals Agency (11) and suggestive of broad activity against all enveloped viruses. *In vitro* exposure of SARS-CoV-2 to chokeberry juice reduced viral titers by more than 96%. This inactivating activity is lower than that observed for oral rinses containing chemicals such as dequalinium chloride and benzalkonium chloride, povidone-iodine, or ethanol and essential oils that show titer reductions of >99.9% (5). Generally, differences in contact times did not have a strong influence on the efficiency of inactivation suggesting a rapid acting antiviral effect. As expected, infectivity of the non-enveloped adenovirus was not or marginally affected by the tested products.

The antiviral activities of the plant products can be based on an acidic pH that may directly inactivate virus particles or by (poly-)phenols such as catechins, tannins or flavonoids that can act on viral and cellular proteins (12, 13). For example, pomegranate polyphenols were shown to inhibit influenza viruses by acting on virion surface glycoproteins and causing structural damage to the virion (14). Similarly, green tea catechins have been shown to destroy the virion structure and *epigallocatechin gallate* aggregates virus particles to prevent their interaction with target cells (15). Catechins not only act on virus particles but have additionally been shown to prevent fusion by interfering with endosome acidification and viral enzymes. Similarly, *in silico* modeling has suggested that theaflavin-3,3’-digallat may prevent SARS-CoV-2 infection by interacting with its cellular receptor angiotensin converting enzyme 2 (ACE2) (13, 15). Of note, the composition of natural food products may vary between batches, which might affect their antiviral efficiency. Nevertheless, the composition of various antivirally active components, acting by different mechanisms, may represent a potent mix interfering with virus infection. However, the underlying antiviral mechanism of the tested products herein remain to be identified.

Since viral replication, symptoms and transmission occur in the naso- and oropharyngeal area, reducing viral titers as early as possible might represent a proactive strategy to prevent infection, dissemination, disease, and spread. The herbal products tested in this *in vitro* study are commonly available food preparations that could be applied as convenient “oral rinses”. Antiseptic oral rinses containing membrane-damaging agents (i.e. ethanol, chlorhexidine, cetylpyridinium chloride, hydrogen peroxide and povidone-iodine) are used in various private or clinical situations for prophylactic and therapeutic purposes and have further been applied in the context of viral infections (4, 5). In contrast to these chemical preparations, green tea and herbal juices can be applied more frequently and may be simply swallowed. Gargling tea, tea extracts or plant juice followed by drinking has already been shown to lower the incidence of influenza virus infections, viral loads and symptoms (8, 9). However, whether similar effects may be observed with antivirally active chokeberry juice against SARS-CoV-2 remains to be addressed in future work. We would like to emphasize that our study was designed to determine the virucidal activity of chosen beverages under experimental *in vitro* conditions that do not reflect the more complex *in vivo* environment of the oral cavity. Thus, well controlled clinical studies are required to clarify whether application of plant juices may reduce viral loads in the oral cavity before any recommendations should be made.

